# Pramipexole restores behavioral inhibition in highly impulsive rats through a paradoxical modulation of frontostriatal networks

**DOI:** 10.1101/2023.02.05.527198

**Authors:** Robin Magnard, Maxime Fouyssac, Yvan M. Vachez, Yifeng Cheng, Thibault Dufourd, Carole Carcenac, Sabrina Boulet, Patricia H. Janak, Marc Savasta, David Belin, Sebastien Carnicella

**Author notes:** **Correspondence:** Dr Robin Magnard; Present address: Department of Psychological and Brain Sciences, Johns Hopkins University, Baltimore, MD, 21218, USA, *Email address:. Co-senior author (David Belin, Sebastien Carnicella).

## Abstract

Impulse control disorders (ICDs), a wide spectrum of maladaptive behaviors which includes pathological gambling, hypersexuality and compulsive buying, have been recently suggested to be triggered or aggravated by treatments with dopamine D_2/3_ receptor agonists, such as pramipexole (PPX). Despite evidence showing that impulsivity is associated with functional alterations in corticostriatal networks, the neural basis of the exacerbation of impulsivity by PPX has not been elucidated. Here we used a hotspot analysis to assess the functional recruitment of several corticostriatal structures by PPX in male rats identified as highly (HI), moderately impulsive (MI) or with low levels of impulsivity (LI) in the 5-choice serial reaction time task (5-CSRTT). PPX dramatically reduced impulsivity in HI rats. Assessment of the expression pattern of the two immediate early genes C-fos and Zif268 by *in situ* hybridization subsequently revealed that PPX resulted in a decrease in Zif268 mRNA levels in different striatal regions of both LI and HI rats accompanied by a high impulsivity specific reduction of Zif268 mRNA levels in prelimbic and cingulate cortices. PPX also decreased C-fos mRNA levels in all striatal regions of LI rats, but only in the dorsolateral striatum and nucleus accumbens core (NAc Core) of HI rats. Structural equation modelling further suggested that the anti-impulsive effect of PPX was mainly attributable to the specific downregulation of Zif268 mRNA in the NAc Core. Altogether, our results show that PPX restores impulse control in highly impulsive rats by modulation of limbic frontostriatal circuits.

## INTRODUCTION

Impulse control disorders (ICDs) represent a set of heterogeneous maladaptive behaviors characterized by a deficit in urge regulation and inhibition that are core symptoms of several neuropsychiatric conditions, that include, for instance, intermittent explosive disorder, pathological gambling and hypersexuality [1].

Despite the profound consequences ICDs have on the quality of life of patients, their neurobiological basis has not yet been elucidated. However ICDs show a relatively high prevalence in Parkinson’s disease (PD) [2,3], restless leg syndrome [4–6], fibromyalgia [7] and hyperprolactinemia [8–10], the treatment of which involves dopamine D_2/3_ receptors agonists such as pramipexole (PPX) or ropinirole. This implicates the iatrogenic effect of dopaminergic drugs and a potential abnormal dopaminergic function in the development of ICDs.

Alongside their apparent reliance on aberrant dopaminergic mechanisms, the compulsive nature of ICDs, which is not without similarities with that of substance use disorders [11–18], has led to their conceptualization as a form of behavioral addiction [19–21]. Similarly to psychostimulant use disorder, the vulnerability to develop ICDs has been shown to be associated with a high impulsivity trait [14,17,21–24], which is characterized by a tendency to act prematurely, without forethought or concerns for adverse upcoming consequences [25]. Impulsivity is a multifaceted construct [26] that encompasses the inability to tolerate delays to reinforcement and adapt to risks, referred to as cognitive impulsivity, on one hand, and the inability to withhold prepotent responses, resulting in poor or adverse outcomes, so-called motor/waiting impulsivity [25], on the other hand. Cognitive impulsivity is principally assessed in delayed discounting [27–29] or risk taking [30–32] tasks whereas waiting impulsivity is canonically assessed as the rate of premature responses in the 5-choice serial reaction time task (5-CSRTT) [33,34].

The respective contribution of each of these dimensions of impulsivity, which are behaviorally and neurally dissociable [17,35–37], to the emergence of ICDs following exposure to dopaminergic drugs such as PPX has not yet been elucidated. While preclinical studies in rodents and non-human primates have demonstrated that PPX exacerbates cognitive impulsivity, as assessed in delay discounting [27–29] and risk taking [30–32] tasks, the influence of PPX on waiting impulsivity is less clear. Indeed, PPX has been shown to exacerbate waiting impulsivity as assessed in differential reinforcement of low rate of responding and fixed consecutive number tasks [38]. In the 5-CSRTT, one study [39] found PPX to have a detrimental effect mainly on accuracy, the index of attention, and to bring omissions rate up to almost 50%, suggesting that the animals were no longer engaged in the task in that study, precluding any assessment of the effect of the drug on waiting impulsivity per se. In addition, the interaction between PPX treatment and impulsivity trait was not investigated.

Considering the heuristic and translational value of high impulsivity trait, characteristic of individuals belonging to the upper quartile of the population ranked on the rate of premature responses in the 5-CSRTT, with regards to the vulnerability to develop compulsive behaviors [40–43], in the present study we sought to test the hypothesis that PPX may exert an impulsivity trait specific effect on impulse control associated with functional alterations in underlying corticostriatal circuits [25]. A pro-impulsive action of PPX in highly impulsive (HI) rats characterized in the 5-CSRTT would provide experimental evidence towards a facilitation of the development of compulsivity by dopaminergic drugs in this vulnerable population.

At the neural systems level, impulsivity has been shown to depend on the functional engagement of a distributed corticostriatal network [25] that involves the insular cortex, the nucleus accumbens core (NAc Core) and shell (NAc Shell) and the dorsomedial striatum (DMS) [25,37,44,45]. In contrast, compulsive behaviors that result from the interaction between a pre-existing impulsive trait and dopaminergic drugs have repeatedly been shown to be associated with, if not mediated by, dopaminergic mechanisms in the dorsolateral striatum (DLS) [46,47]. Thus, using *in situ* hybridization we quantified the mRNA levels of the cellular activity and plasticity markers C-fos [48–50] and Zif268 [51–55], respectively, in corticostriatal structures and used a mediation analysis that aimed to identify the functional signature of the effect of PPX on impulsivity in vulnerable individuals.

## MATERIALS AND METHODS

### Subjects

Experiments were performed on 48 male Sprague-Dawley rats (Janvier, France) that were 6 weeks old (weighing 200 g) when they were habituated to the animal facility in which they were housed 1 per cage under a 12 h/light/dark cycle (lights ON at 7 am). Rats were food restricted to 90% of their theoretical free feeding weight for 5-CSRTT training, but had *ad libitum* access to water throughout the experiment. Protocols complied with the European Union 2010 Animal Welfare Act and the new French directive 2010/63 and were approved by the French national ethics committee n°004.

### 5-CSRTT Training

The 5-CSRTT procedure has been adapted for Sprague-Dawley from previous studies [33,56–59]. Briefly, the apparatus consisted of eight 25 x 25 x 25 cm operant chambers (Med associates, St Albans, VT) each equipped with a curved rear wall. Set into the curved walls were five 2.5 x 2.5 cm square holes, 4 cm deep and 2 cm above the floor. Each hole was equipped with an infra-red beam crossing the entrance horizontally and a cue light at its rear that provided illumination. A houselight was located at the top of this wall. 45 mg sucrose pellets (TestDiet, VA) were delivered from a pellet dispenser to a tray at the front of the cage, also equipped with a cue light and infra-red beam.

#### Phase 1: Learning

The procedure began one week after food restriction was initiated. To prevent neophobia, rats were exposed to 10 sucrose pellets in their home cage for two consecutive days. The following two days, rats were placed in their operant box for 15 min habituation sessions, during which ten pellets were placed in the magazine and two pellets were placed in each hole, all cue- and house-lights were turned ON.

#### Phase 2: Task acquisition

As detailed in **Fig. 1A** each daily session consisted of 100 discrete trials with stable performance being achieved after about 40 sessions. In order to facilitate brief stimulus detection by Sprague-Dawley rats, the house-light remained OFF during the length of the session and was turned on during time out periods [60]. Rats were trained to enter the food magazine to initiate a trial. After a 5 s intertrial interval (ITI), a brief light stimulus was pseudo-randomly presented in one of five holes to indicate the individual the location of the correct response for that trial. Following a nosepoke in this hole (‘correct response’), rats were rewarded with the delivery of one sucrose pellet in the food tray. A nosepoke response in any of the adjacent holes (‘incorrect response’), or a failure to respond within 5 s after the onset of the stimulus (‘omission’), resulted in no pellet delivery and a 5 s time-out period signaled by the house light being turned ON. Additional nosepokes performed after a correct or incorrect response (referred as ‘perseverative response’) were recorded but had no consequence. Nosepokes made during the ITI, that is, before the onset of the stimulus (or ‘premature responses’) were recorded as a measure of waiting impulsivity, and resulted in a 5 s time-out (house light turned ON) and reward omission. Nosepokes made during this time out period (‘time-out response’) were recorded, but had no programed consequence. The stimulus duration was progressively reduced from 20 s to 1 s across training stages, as previously described [58,59]. Progress across the different stages was determined by at least 80% accuracy and less than 20% omissions on any particular stage. Rats which failed to fulfil these criteria were excluded from the study (n = 4).

**Figure 1:**
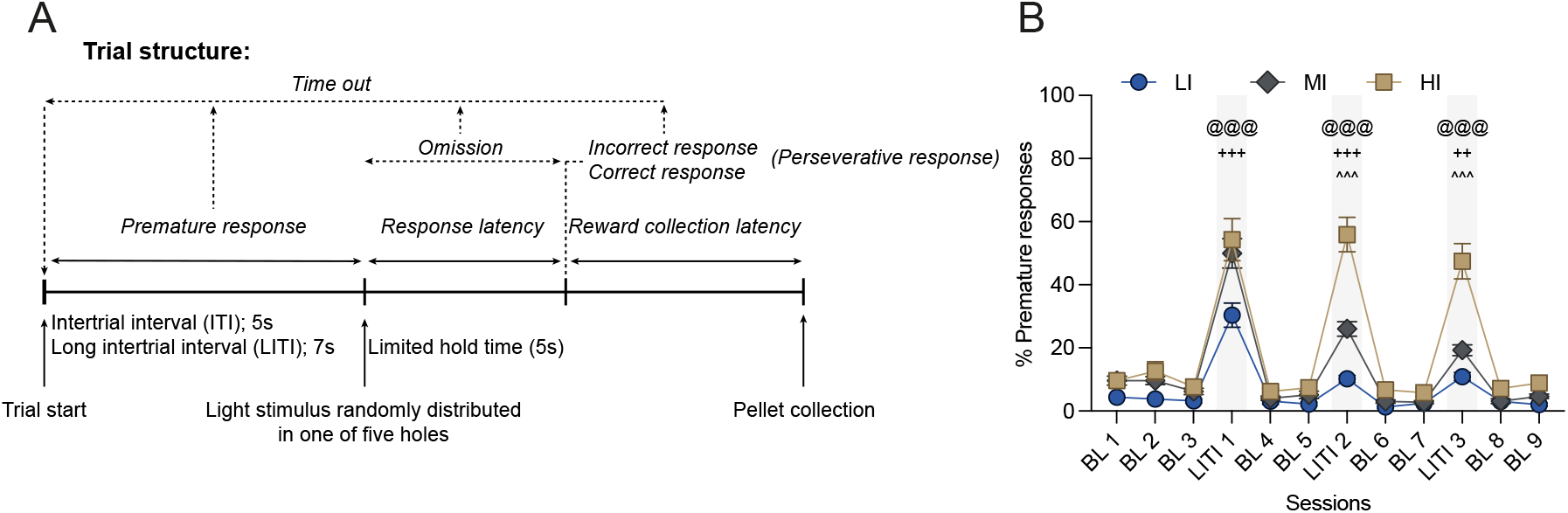
Structure of the 5-CSRTT and identification of low, moderate and high impulsivity rats. (**A**) 5-CSRTT trial structure. (**B**) Segregation of low impulsive (LI, n = 11), medium (MI, n = 22) and highly impulsive (HI, n = 11) rats. BL: baseline; LITI: long intertrial interval. Data are shown as means ± SEM. HI vs. LI ^@@@^p<0.001; HI vs. MI ^^^p<0.001; MI vs. LI ^++^p<0.01; ^+++^p<0.001.

#### Phase 3: Challenge with long intertrial interval

Following acquisition of the task, rats were challenged with three 7 s-long intertrial interval (LITI) sessions separated by two baseline sessions with the regular 5 s ITI. Subjects were ranked according to their level of premature responses during second and third LITI [45] and those in the upper and lower quartile (n = 11 each) [45] were considered as having high (HI) or low impulsivity (LI), respectively. The rest of the population was considered to be moderately impulsive (MI, n = 22).

### Pharmacological procedures

Individuals of the HI, MI and LI groups were randomly allocated to a PPX or a vehicle treatment group so that the PPX-treated HI and LI rats (n = 6 and 5 respectively), as well as MI rats (n = 12) showed a similar level of premature responses before initiation of treatment as that shown by Veh-treated HI, LI and MI rats (n = 6, n = 5 and n = 10, respectively). Rats received an intraperitoneal administration of 0.2 mg/kg/day PPX (Sigma–Aldrich, St. Louis, MO), diluted in 0.9% NaCl, or vehicle (Veh) 30 minutes prior to each daily behavioral session, with treatment starting one day before to the first of twelve sessions (11 baseline 5 s ITI sessions and one final 7 s LITI session) over which the effect PPX on impulsivity in HI and LI rats was measured. The dose of pramipexole used here was carefully chosen because it exacerbates impulsivity in a delay discounting task (Magnard & Carnicella, unpublished) and improve motivational function in dopamine-deficient rats [61] (see for review [62]), while it does not influence locomotion [63,64].

### Tissue collection

15 minutes after the final LITI session, rats were deeply anesthetized by isoflurane and killed by decapitation. Brains were quickly removed and snap frozen in liquid nitrogen, then stored at -80°C until they were processed into 14μm-thick coronal brain sections with a cryostat (Microm HM 500, Microm, Francheville, France) and collected on permafrost gelatin-coated slides (Colorfrost Plus, Fisher scientific, Pittsburg, PA) that were stored at -80°C.

### In situ Hybridization

The *in situ* hybridization (ISH) procedure was carried out as previously described [65,66] with oligonucleotide probes specifically complementary to the sequence of the mRNA of the immediate early genes C-Fos (nucleotides 159-203 of the NCBI Reference Sequence NM_022197.2) or Zif-268 (nucleotides 1680-1724 of the NCBI Reference Sequence NM_012551.3), tailed by 3’OH incorporation of ^35^S-dATP (1250 mCi/mmol, Perkin Elmer, UK) by a terminal Deoxynucleotidyl transferase (Promega, M1875) with a specificity of 2.5 x 10^6^ cpm/ml (C-Fos) or 3 x 10^6^ cpm/ml (Zif-268).

Following fixation and pre-hybridization treatments aiming at reducing non-specific hybridization, slides were incubated overnight at 42ºC in the hybridization buffer [*50% deionized formamide, 10% dextran sulfate, 50ng/ml denaturated salmon sperm DNA, 5% Sarcosyl, 0*.*2% SDS, 1mM EDTA, 300nM NaCl, 5X Denhardt’s in 2X standard sodium citrate (SSC)*] with the probes diluted at a concentration of 8.75ng/ml (C-Fos) or 6.25ng/ml (Zif-268). Slides were then washed in decreasing concentration of SSC and dehydrated in increased ethanol concentration baths. Sections were exposed to Biomax MR films (Kodak, Rochester, USA) for four weeks (Zif268-labelled sections) or 6 weeks (C-Fos-labelled sections) at room temperature. Films were revealed in a dark room. Pictures of each brain section were taken on a Northern light (Imaging Res Inc.) light table with a Qicam (QImaging) camera equipped with a SIGMA 50 mm 1:2.8 DG MacroD Fast 1394 (Nikon) objective and subsequently analyzed with ImageJ software [67]. A region of interest was drawn for each striatal territory in which the optical density reflective of the mRNA level was measured (according to the rat brain atlas [68]). The optical density in an mRNA-free part of the brain (i.e., corpus callosum) was defined as background, and this value was subtracted to that obtained from the area of interest to compute the relative optical density used as the dependent variable in subsequent analyses.

### Data and statistical analyses

Data, presented as mean ± SEM with or without superimposed individual data points, were analyzed using SigmaStat (Systat software Inc., San Jose, USA) and SPSS (IBM, Amorak, NY). Three-way repeated measure analyses of variance (RM-ANOVAs) were used to compare groups across sessions, with sessions as within-subject factor, impulsivity (LI, MI or HI) and treatment (Veh or PPX) as between-subject factors. Two-way RM ANOVAs were used to compare levels of impulsivity between LI, MI and HI rats during the screening period, with sessions as within-subject factor and impulsivity as between-subject factor. Two-way ANOVAs were also used to compare LI, MI and HI rats for BL11 and LITI sessions with impulsivity and treatment as between-factors. Two-way ANCOVA was used to control for the omission and BL11 premature response performances on premature responses during LITI, with impulsivity and treatment as the between-subject variables, LITI omissions and BL11 premature responses as covariants.

Differences between LI, MI and HI rats in Zif268 and C-fos mRNA were analyzed using three-way ANOVA with impulsivity and treatment as between-subject factors and structures as within-subject factor. When indicated, *post hoc* analyses were carried-out using the Student-Newman-Keuls test.

Dimensional inter-relationships were analyzed with nonparametric Spearman correlation coefficient ρ and Spearman rank order tests. Assumptions for the normality of the distributions and the homogeneity of variance were verified using the Shapiro-Wilk and Levene test, respectively. Significant violations of homogeneity of variances and normality were corrected using square root transformations. Significance for p values was set at α = 0.05. Effect sizes for the ANOVAs are also reported using partial η^2^ values (η_p_^2^) [69,70].

In order better to delineate the neural basis of the influence of PPX on impulsivity, we employed mediation analysis [71,72] that helped decipher the direct effect of treatment and the indirect effect mediated by C-fos and Zif268 expression on impulsivity. For this, we applied:

a. dependent regression (direct effect)

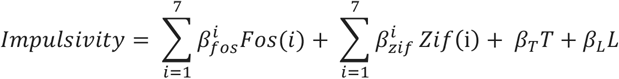
b. mediator regression

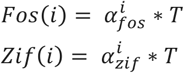
c. effect decomposition (indirect effect)

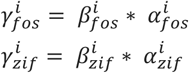

where *Fos(i)*and *zif(i)*represent the mRNA levels of the immediate early genes (IEG) C-fos and Zif268, respectively, from each brain region. *i* represents the arbitrary index of the seven brain regions we stained (IL, PrL, Cg, DLS, DMS, NAc Core, NAc Shell). *T* is a binary variable (1 for PPX, 0 for Veh). *L* is impulsivity measured under LITI prior to pharmacological treatment. β represents the direct effect from each variable. α represents the mediation effect from treatment (T) to cellular activity/plasticity. γ represents the indirect effect of treatment on impulsivity mediated by the two IEGs. To strengthen the analysis and prevent any bias by analyzing only a subpopulation of this study, we included LI, MI and HI -Veh and PPX-treated animals to this analysis which was carried out using the JASP statistical software with bootstrap method, 1000 replications and bias-corrected percentile [71,72].

## RESULTS

### 5-CSRTT screening for low impulsive and high impulsive rats

As previously described [37,45] when Sprague Dawley rats well-trained in a 5-CSRTT are challenged with longer ITIs, thereby requiring individuals to refrain from expressing prepotent responses for a slightly longer period of time than they usual (**Fig. 1A**), marked individual differences in waiting impulsivity are revealed that allow identification of HI and LI rats in the upper and lower quartiles of the population, respectively, and MI in the two middle quartiles. In accordance, HI rats displayed a greater increase in premature responses than LI rats during the LITI sessions (**Fig. 1B**) [impulsivity x session interaction: F_22,451_ = 11.74, p < 0.001, η_p_^2^ = 0.36].

### PPX influences impulse control differentially in LI and HI rats

LI, MI and HI rats were then split into two treatment groups that did not differ in their baseline impulsivity level [no impulsivity x pretreatment x session interaction: F_22,418_ < 1, p = 0.914, η_p_^2^ = 0.03]. Treatment groups received daily IP injections of either PPX (LI, MI or HI PPX-treated rats) or vehicle (LI, MI or HI Veh-treated rats) for 12 sessions.

PPX treatment was revealed to influence differentially baseline impulsivity and its exacerbation by LITI mostly in HI rats (**Fig. 2A**) [sessions x impulsivity x treatment interaction: F_22,418_ = 2.72, p < 0.001, η_p_^2^ = 0.125]. While HI-Veh rats did not differ from LI-Veh rats during baseline sessions, PPX treatment resulted in a progressive increase in baseline premature responses only in HI rats (**Fig. 2A and B**, BL white panels). In marked contrast, PPX treatment prevented the exacerbation of premature responses otherwise shown by HI-Veh and MI-Veh rats upon introduction of LITI (**Fig. 2A and B**, LITI grey panels) [impulsivity x treatment interaction: F_2,44_ = 3.75, p = 0.03, η_p_^2^ = 0.16].

**Figure 2:**
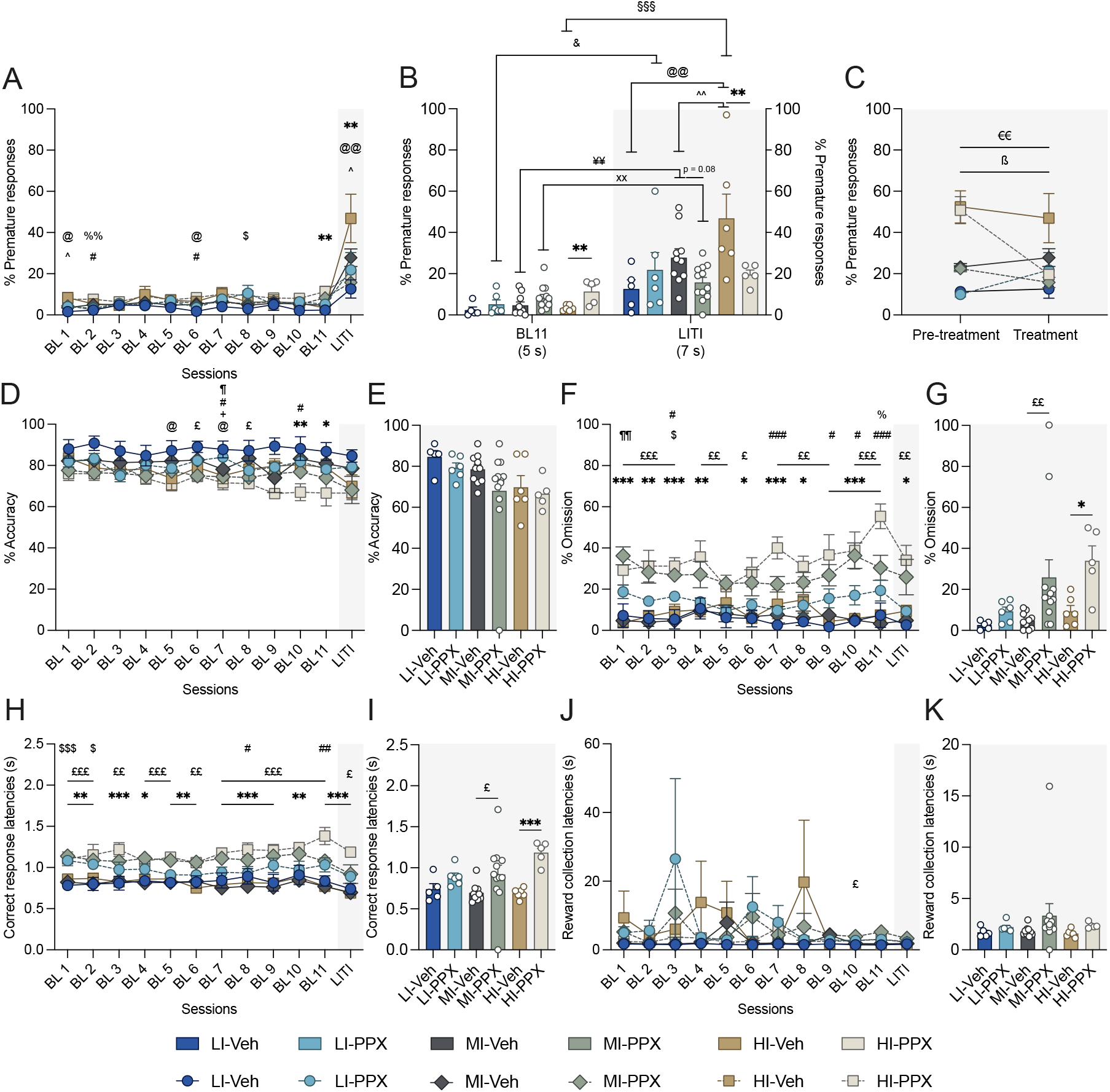
PPX influences premature responses differentially in LI, MI and HI rats in baseline and LITI conditions. (**A**) Percentage of premature responses across baseline (BL) and long intertrial interval (LITI) sessions. (**B**) Differential effect of PPX on premature responses between the last BL and the LITI session. (**C**) Comparison of the level of premature responding during LITI before and during differential PPX vs. vehicle (Veh) treatment. (**D**) Response accuracy across BL and LITI sessions. (**E**) Response accuracy during the LITI session under differential PPX vs. Veh treatment. (**F**) Response omissions across BL and LITI sessions. (**G**) Response omission during the LITI session under differential PPX vs. Veh treatment. (**H**) Correct response latencies across BL and LITI sessions. (**I**) Correct response latency during the LITI session under differential PPX vs. Veh treatment. (**J**) Reward collection latency across BL and LITI sessions. (**K**) Reward collection latency during the LITI session under differential PPX vs. Veh treatment. LI-Veh n= 5; LI-PPX n = 6; MI-Veh n = 10; MI-PPX n = 12; HI-Veh n = 6; HI-PPX n = 5. LI: low impulsive; Moderately impulsive; HI: highly impulsive. Data are shown as means ± SEM. HI-Veh vs. LI-Veh ^@^p<0.05; ^@@^p<0.01. HI-Veh vs. MI-Veh ^^^p<0.05; ^^^^p<0.01. MI-Veh vs LI-Veh ^+^p<0.05. LI-Veh vs. LI-PPX ^$^p<0.05; ^$$$^p<0.001. MI-Veh vs MI-PPX ^£^p<0.05; ^££^p<0.01; ^£££^p<0.001. HI-Veh vs HI-PPX ^*^p<0.05; ^**^p<0.01; ^***^p<0.001. HI-PPX vs. MI-PPX ^%^p<0.05; ^%%^p<0.01. HI-PPX vs. LI-PPX ^#^p<0.05; ^##^p<0.01; ^###^p<0.001. LI-PPX vs. MI-PPX p<0.05; ^¶¶^p<0.01. LI-PPX BL11 vs. LI-PPX LITI ^&^p<0.05. HI-Veh BL11 vs. HI-Veh LITI ^§§§^<0.001. MI-Veh BL11 vs. MI-Veh LITI ^¥¥^<0.01. MI-PPX BL11 vs. MI-PPX LITI ^xx^<0.01. HI-PPX prior treatment vs. HI-PPX under treatment ^€€^p<0.01. MI-PPX prior treatment vs. MI-PPX under treatment ^ß^p<0.05.

This effect was not due to a difference in baseline levels of impulsivity as it persisted when the premature responses expressed during the baseline session that preceded the LITI (BL11) was used as a covariate in an ANCOVA [impulsivity x treatment interaction: F_2,44_ = 6.4, p = 0.004, η_p_^2^ = 0.257]. Within-subject analyses across LITIs confirmed that PPX decreased the level of premature responding under LITI as compared to pre-treatment performance in HI and to a lesser extent in MI rats (**Fig. 2C**) [period x impulsivity x treatment interaction: F_2,38_ = 4.11, p = 0.024, η_p_^2^ = 0.178]. Thus, while PPX treatment moderately increased impulsivity in HI rats in baseline conditions, it abolished its exacerbation during LITI sessions, when it is most challenged and likely under greater influenced of negative urgency.

### The effects of PPX were specific to impulsivity

PPX treated rats showed a progressive decline in accuracy over the course of treatment (**Fig. 2D**) [treatment: F_1,38_ = 10.27, p = 0.003, η_p_^2^ = 0.21], which may reflect an impairment in attention [34]. However, this effect was not dependent on impulsive trait [no sessions x impulsivity interaction F_22,418_ < 1, p = 0.57, η_p_^2^ = 0.05; no sessions x impulsivity x treatment interaction F_22,418_ = 1.02, p = 0.44, η_p_^2^ = 0.05]. This result was however not supported by the analysis of performance during the LITI session (**Fig. 2E**) [treatment: F_2,44_ = 1.9, p = 0.17, η_p_^2^ = 0.05; no treatment x impulsivity interaction: F_2,44_ = 0.27, p = 0.76, η_p_^2^ = 0.014]. Together these results demonstrate that under the present experimental conditions, a moderate daily dose of PPX, had a restricted effect on attention, that was not dependent on impulsive phenotype.

In addition, we assessed response omissions and collection latencies, which jointly not only provide an insight into task engagement [34] but could also potentially influence the rate of premature responses. PPX increased the percentage of omitted trials (**Fig. 2F**) [sessions x treatment interaction F_11,418_ = 2.5, p = 0.005, η_p_^2^ = 0.06; no sessions x impulsivity x treatment interaction F_22,418_ = 0.67, p = 0.86, η_p_^2^ = 0.03]. Because this effect was exacerbated in MI and HI rats irrespective of ITI duration [impulsivity x treatment interaction F_2,38_ = 3.26, p = 0.049, η_p_^2^ = 0.147], even though only an effect of the treatment was observed during the LITI session (**Fig 2G**) [treatment: F_1,44_ = 10, p = 0.003, η_p_^2^ = 0.20; no impulsivity x treatment interaction F_2,44_ < 1, p = 0.43, η_p_^2^ = 0.043] an ANCOVA was carried out to determine a potential influence of this PPX-induced increase in omissions on impulsivity using omissions expressed during the LITI session as a covariate. This analysis revealed that the effect of PPX on omissions during the LITI session cannot account for its profound effect on premature responses [impulsivity x treatment interaction: F_2,44_ = 4.15, p = 0.024, η_p_^2^ = 0.187]. Whereas PPX also increased the latencies to correct responses irrespective of the impulsive phenotype, (**Fig. 2H**) [treatment F_1,38_ = 45.5, p < 0.001, η_p_^2^ = 0.54; no sessions x impulsivity x treatment interaction: F_22,418_ = 1,37, p = 0.18, η_p_^2^ = 0.06; no impulsivity x treatment interaction F_2,38_ = 2.14, p = 0.13, η_p_^2^ = 0.10] and (**Fig. 2I**) [no impulsivity x treatment interaction: F_2,44_ = 1.78, p = 0.18, η_p_^2^ = 0.08; treatment F_1,44_ = 15.42, p < 0.001, η_p_^2^ = 0.28], it did not influence reward collection latencies (**Fig. 2J**) [no sessions x impulsivity x treatment interaction, F_22,418_ < 1, p = 0.85, η_p_^2^ = 0.035; treatment F_1,38_ < 1, p = 0.42, η_p_^2^ = 0.017] and (**Fig. 2K**) [treatment: F_1,44_ = 1.7, p = 0.19, η_p_^2^ = 0.04; no impulsivity x treatment interaction: F_2,44_ < 1, p = 0.79, η_p_^2^ = 0.01].

Thus, the slight decrease in response readiness caused by PPX especially in HI rats, is unlikely to account for the effect of this dopaminergic drug on impulsivity. This was further supported by the absence of correlation between premature responses performed during the final LITI session, under differential treatment, and the response rate before treatment during either baseline (**Suppl. Fig. 1A**) [ρ = -0.133, p = 0.391] (see also **Suppl. Fig. 1B and C** for LI, MI and HI rats, respectively) or LITI sessions (**Suppl. Fig. 1D**) [ρ = 0.184, p = 0.233] (see also **Suppl. Fig. 1E** and **F** for LI, MI and HI rats, respectively), thus confirming that the effect of PPX on response readiness did not influence premature responses.

Finally, PPX did not influence perseverative responses (**Suppl. Fig. 2A**) [no sessions x impulsivity x treatment interaction, F_22,418_ < 1, p = 0.92, η_p_^2^ = 0.03; no impulsivity x treatment interaction F_3,38_ < 1, p = 0.63, η_p_^2^ = 0.02], but reduced magazine entries reminiscent of reduced response readiness noticed in PPX treated rats (**Suppl. Fig. 2B**) [session x treatment interaction: F_11,418_ = 2.9, p < 0.001, η_p_^2^ = 0.07 no sessions x impulsivity x treatment interaction, F_22,418_ < 1, p = 0.82, η_p_^2^ = 0.03; no impulsivity x treatment interaction: F_2,38_ = 1.3, p = 0.26, η_p_^2^ = 0.06]. Therefore, PPX treatment did not induce repetitive or stereotyped behaviors that may influence performance in the 5-CSRTT.

### The dampening effect of PPX on the exacerbated impulsivity during LITI sessions in HI rats was associated with a functional engagement of corticostriatal circuits

In order to identify the neural basis of the effect of PPX on impulsivity trait, we carried out a hot-spot analysis in structures of the corticostriatal circuitry using *in situ* hybridization for the markers of cellular activity and plasticity C-fos and Zif268, respectively.

A first quality control analysis of the pattern of expression of the two IEGs, based on a covariance analysis for each marker, revealed a convergence of RNA levels in line with the functional organization of the corticostriatal circuitry, with similar trends in expression in structures inter-connected to each other (**Suppl. Fig. 3**). However, as anticipated, the two IEGs were shown here not to reflect the same functional process as their mRNA levels did not covary (**Suppl. Fig. 3**).

**Figure 3:**
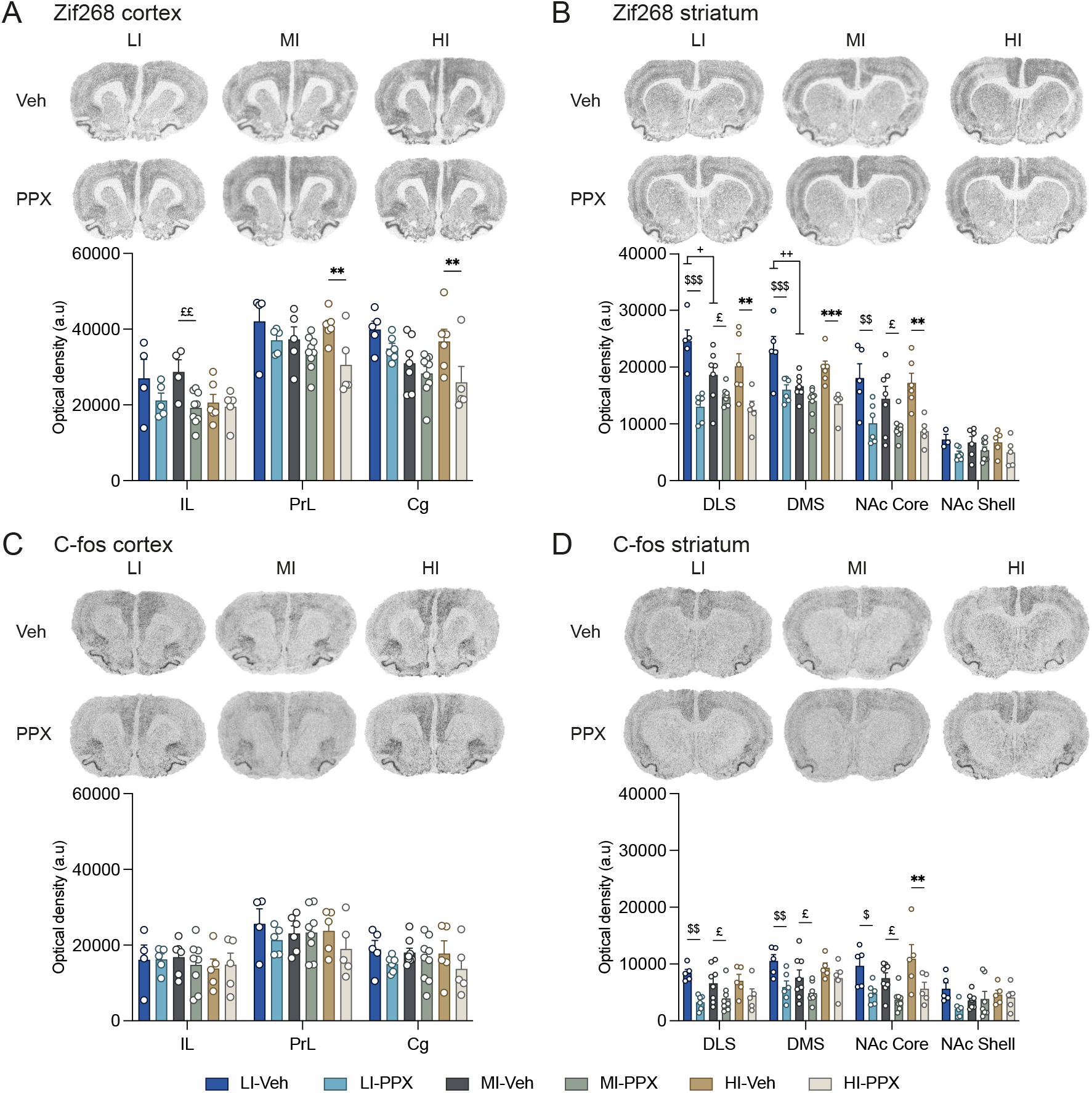
Zif268- and C-fos-based functional signature of the effect of PPX on impulsivity in the prefrontal corticostriatal circuitry of LI, MI and HI rats. (**A**) Zif268 mRNA levels quantified in cortical areas. Upper panel: representative high-resolution photos of the signal obtained following in situ hybridization with a probe against Zif268 in Veh- and PPX-treated LI, MI and HI rats. Lower panel: Optical densities measured in the IL, the PrL and the Cg cortex. (**B**) Zif268 mRNA levels quantified in the striatum. Upper panel: representative high-resolution photos of the signal obtained following in situ hybridization with a probe against Zif268 in Veh- and PPX-treated LI, MI and HI rats. Lower panel: Optical density measured in the DLS, the DMS, the NAc Core and the NAc Shell. (**C**) C-fos mRNA levels quantified in cortical areas. Upper panel: representative high-resolution photos of the signal obtained following in situ hybridization with a probe against C-fos in Veh- and PPX-treated LI, MI and HI rats. Lower panel: Optical densities measured in the IL, the PrL and the Cg cortex. (**D**) C-fos mRNA levels quantified in the striatum. Upper panel: representative high-resolution photos of the signal obtained following in situ hybridization with a probe against C-fos in Veh- and PPX-treated LI, MI and HI rats. Lower panel: Optical density measured in the DLS, the DMS, the NAc Core and the NAc Shell. Photos per different condition represent the same animal stained with different in situ hybridization probe. LI-Veh n= 4-5; LI-PPX n = 5-6; MI-Veh n = 4-8; MI-PPX n = 7-9; HI-Veh n = 5-6; HI-PPX n = 5. LI: low impulsive; MI: moderately impulsive; HI: highly impulsive; IL: infralimbic cortex; PrL: prelimbic cortex; Cg: cingulate cortex; DLS: dorsolateral striatum; DMS: dorsomedial striatum; NAc Core: nucleus accumbens core; NAc Shell: nucleus accumbens shell. Data are shown as means ± SEM. HI-PPX vs. HI-Veh ^*^p<0.05; ^**^p<0.01. HI-PPX vs. LI-PPX ^#^p<0.05. LI-Veh vs. LI-PPX ^$^p<0.05; ^$$^p<0.01; ^$$$^p<0.001. MI-Veh vs. MI-PPX ^£^p<0.05; ^££^p<0.01. LI-Veh vs. MI-Veh ^+^p<0.05; ^++^p<0.01.

An impulsive trait dependent reduction of Zif268 was observed in the dorsal structures of the mPFC, namely the prelimbic and cingulate cortexes of PPX treated rats (**Fig. 3A**) [structure x impulsivity x treatment interaction: F_4,54_ = 3.18, p = 0.02, η_p_^2^ = 0.19].

This decreased recruitment of a marker of cellular plasticity in the prefrontal cortical areas was accompanied by a decrease in the level of expression of Zif268 across the ventral and dorsal domains of the striatum in PPX-treated rats (**Fig 3B**) [structure x treatment interaction: F_3,84_ = 11.12, p < 0.001, η_p_^2^ = 0.28; no structure x impulsivity x treatment interaction: F_6,84_ = 0.53, p = 0.78, η_p_^2^ = 0.037]. While the decrease in Zif268 mRNA levels in the DLS was quantitatively similar to that observed in the DMS, differences were observed between the NAc Core and NAc Shell in that Zif268 mRNA levels were much more reduced by PPX in the Core than the Shell.

The effect of PPX on the levels of C-fos mRNA in the prefrontal regions of LI, MI and HI rats followed the same trend as that observed for Zif268, albeit of a lesser magnitude (**Fig. 3A**) [Structure x impulsivity x treatment interaction: F_4,54_ = 2.77, p = 0.036, η_p_^2^ = 0.17]. However, structure level analysis failed to isolate significant differences between the different treatment or impulsivity conditions.

In the striatum, PPX treatment resulted in a global decrease in C-fos mRNA levels in the DLS, DMS, NAc Core, but no difference was observed in the NAc Shell (**Fig. 3B)** [Structure x treatment interaction: F_3,90_ = 4.24, p = 0.007, η_p_^2^ = 0.12]. This effect of the treatment was found to be independent of the impulsive phenotype of these rats [no structure x impulsivity x treatment interaction: F_6,90_ < 1, p = 0.71, η_p_^2^ = 0.04]. Together these results suggest that a high impulsivity trait dampens the inhibition exerted by PPX on NAc Shell and DMS activity, as assessed by C-fos mRNA levels.

### PPX-induced alterations of Zif268 mRNA levels in a network involving the PrL, the NAc Core and the DLS predicted the anti-impulsive effect of the drug

We then carried-out a mediation analysis in order to identify which of the functional changes in the prefrontal corticostriatal circuitry described above predicted best the anti-impulsive effect of PPX. Mediation analysis tests the validity of a hypothetical causal chain in which one variable, here PPX treatment, affects a second variable, here Zif268 and C-fos expression, which, in turn, affects a third variable, impulsivity, in a so-called indirect effect. Thus, in our statistical model, mRNA levels of Zif268 or C-fos in the different corticostriatal structures would “mediate” the relationship between the predictor, treatment (PPX), and the behavioral outcome, impulsivity (premature responses during the last LITI under treatment) (**Fig. 4A and B**). The first outcome of the analysis was a negative estimate c between treatment and impulsivity (**Fig. 4C** upper panel) which confirmed the reduction of LITI-exacerbated impulsivity by PPX, thereby representing a first validation of the model. The indirect effect analysis also confirmed that PPX decreased both C-fos and Zif268 mRNA expression in the seven structures investigated (i.e negative estimate a; **Fig. 4C** lower panel). While no significant relationship with C-fos expression was identified, the mediation analysis revealed that a reduction of Zif268 mRNA level in the NAc Core, PrL and DLS was correlated with PPX-induced changes in premature responding. More specifically, at the cohort level, a reduction of Zif268 expression in the NAc Core was positively correlated to a reduction in impulsivity (i.e positive estimate b) whereas in the PrL and the DLS it was associated with increased impulsivity (i.e negative estimate b). Overall, these results enabled the identification of a network involving the PrL, NAc Core and DLS as the neural substrate of the anti-impulsive effects of PPX.

**Figure 4:**
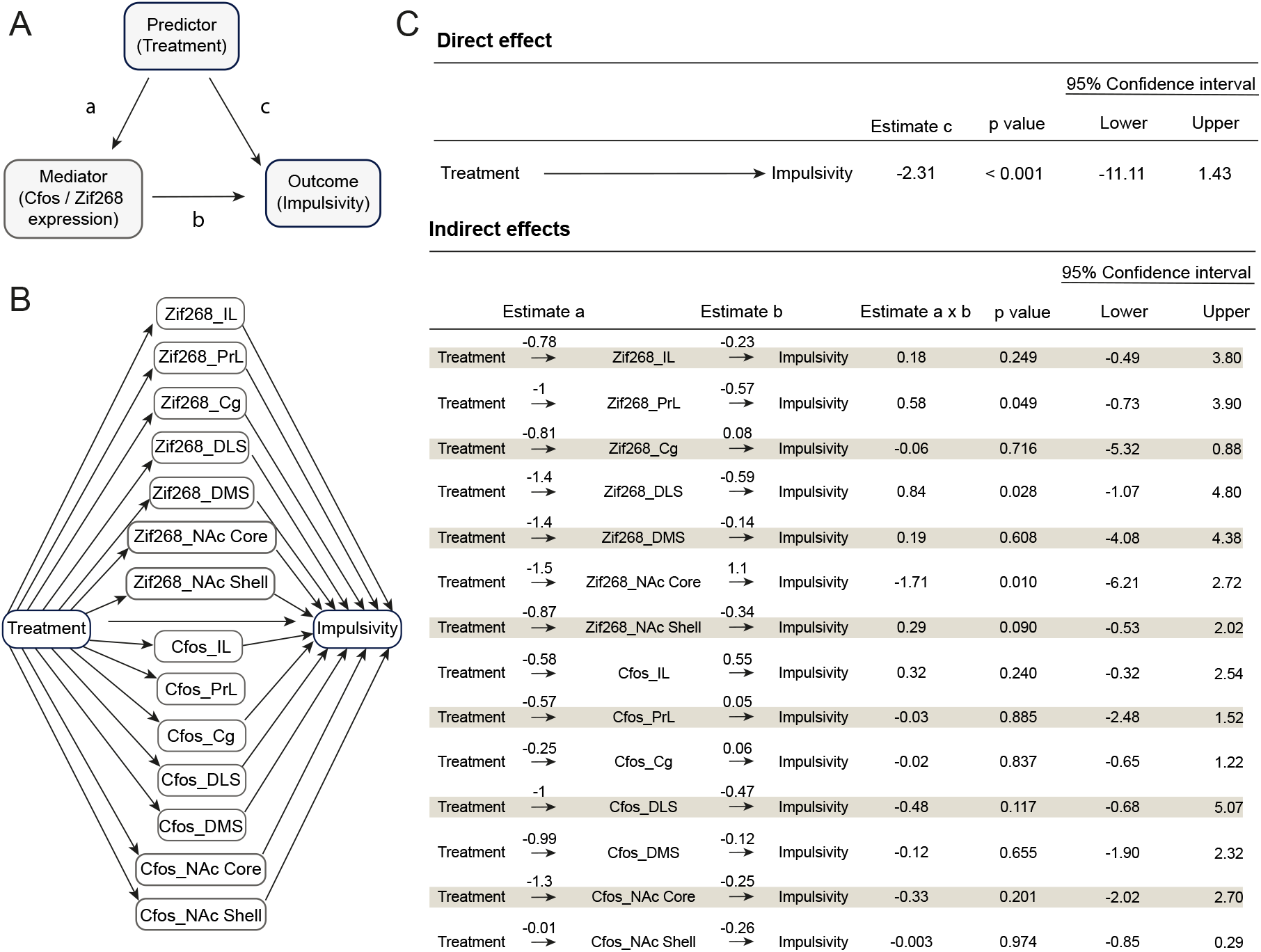
The PPX-induced alterations of Zif268 mRNA levels in a network involving the PrL, the NAc Core and the DLS predicted the anti-impulsive effect of the drug. (**A-B**) Simplified diagrams of the mediation analysis. (**C**) Statistical results of the mediation analysis. Upper panel: direct effect of the treatment on premature responses (i.e., impulsivity). Lower panel: indirect effects of the treatment on premature responses mediated by IEGs mRNA levels. LI-Veh n= 5; LI-PPX n = 6; MI-Veh n = 8; MI-PPX n = 9; HI-Veh n = 6; HI-PPX n = 5. LI: low impulsive; MI: moderately impulsive; HI: highly impulsive.

## DISCUSSION

This study provides a detailed behavioral characterization of the effect of PPX on the expression of a high impulsivity trait either at baseline or when challenged under long inter-trial interval conditions. Repeated PPX administration was shown to exacerbate baseline impulsivity, assessed across several 5 s ITI sessions, only in HI rats. In marked contrast, PPX prevented the exacerbation of premature responding characteristic of HI and MI rats, to a lesser extent, upon introduction of a longer 7s ITI. In addition, the gene expression hotspot analyses carried out on the entire population revealed that the trait-dependent effect of PPX on impulse control is mediated by functional alterations of a corticostriatal network involving the PrL, the NAc Core and the DLS.

These results together challenge an established wisdom about the pro-impulsivity effects of D_2_-like agonists since they are demonstrated here to be dependent on pre-existing differences in impulsivity. Impulsivity trait-dependent effects have already been described for the noradrenaline reuptake inhibitor, atomoxetine, and the psychostimulant drugs, amphetamine and methylphenidate, which all decrease premature responding under LITI in HI rats but potentiate that of LI rats [42,74–76] (but see [77]). NAc Core deep brain stimulation (DBS) has been shown to exert the same impulsivity trait-dependent effect on impulse control in the 5-CSRTT by reducing premature responses in HI rats and tending to increase them in LI rats. Together these data highlight profound neurobiological differences between HI and LI individuals [78], which are in part associated with a reduced expression of D_2/3_R in the ventral striatum of HI individuals, as shown by two PET scan and ISH studies [45,75,79]. HI rats have also been shown to lower D_2_ mRNA levels both in the nucleus accumbens and the ventral tegmental area than LI rats, which suggests that their lower accumbal D_2_ dopamine receptor level, as assessed in PET, is attributable both to a pre and a post-synaptic decrease. Together these data suggest that the impulsivity trait-dependent effect of PPX cannot be accounted for by a differential engagement of pre-vs post-synaptic striatal D2 dopamine receptors.

The effect of PPX on baseline impulsivity in HI rats observed in the present study is in agreement with the previous demonstration that PPX exacerbates waiting impulsivity in non-parkinsonian animals in a DRL task [38]. In another study that used a 5-CSRTT similar to the one used here, PPX was shown to promote premature responses only in animals with a virus-mediated α-synuclein-induced nigrostriatal lesion. This effect that was exacerbated during LITI, but it was not restricted to HI rats, and was instead accompanied with attentional and motivational deficits which precluded any interpretation with regards to the specific effects of PPX on impulsivity [39]. In contrast, the only side effect of the PPX treatment observed in the present study was a reduction in general activity (see [61]) or motor readiness [73] since deficits in attention or disengagement from the task were not observed in PPX-treated animals.

In marked contrast to its effect on baseline impulsivity, PPX completely blocked the exacerbation of impulsivity by an acute increase in waiting time, as operationalized under LITI, in HI rats. This observation highlights the influence of the engagement of negative urgency in highly impulsive individuals as a factor that may contribute to the switch from a pro- to an anti-impulsive effect of these drugs. Indeed, the increase in waiting time during LITI profoundly challenges the inhibitory control of highly impulsive individuals [80,81] which experience more often the negative consequences of their premature responses, a loss in reinforcement that has been suggested to generate heightened negative urgency [82]. In a recent study, such negative urgency, the trait of making a harsh decision under stress, has been shown to mediate compulsive goal-directed relapse in individuals having developed with a DLS dopamine-dependent incentive habit for cocaine that had lost the opportunity to express it [66].

In line with this interpretation, the ITI-dependent effect of PPX on impulsive control in HI rats observed in this study may reflect the differential effect of dopaminergic drugs on the corticostriatal networks involved in habitual vs. aberrant goal-directed control over behavior in HI rats. Thus, following the overtraining that is inherent to the acquisition of the 5-CSRTT, daily stimulation of the D_2/3_ receptors by PPX during baseline conditions may interfere with instrumental responding mediated by a 5 s timing-based DLS-dependent stimulus-response association [83,84], thereby resulting in a progressive increase in premature responses in HI rats over time. However, during LITI sessions, the changes in timing-dependent action control and heightened negative urgency are likely to result in the engagement of a new PrL/Cg-DMS-dependent action-outcome process, the functional engagement of which is necessary for inhibitory control [85,86]. Under the LITI condition PPX resulted in a selective decrease in C-fos mRNA levels in the NAc Core and the DLS of HI rats, in the absence of any change in the cellular activity in the prefrontal regions. Therefore, the sustained functional engagement of the circuit involving the mPFC and the DMS alongside a decreased functional recruitment of the DLS observed in HI-PPX rats, is strongly suggestive of the engagement of the goal-directed system [25,85].

Hot-spot mapping was carried out in parallel for C-fos and Zif268 because while these two immediate early genes have long been shown to be recruited during cellular activity and plasticity events, their recruitment is influenced by different cellular mechanisms and follows distinct, partially overlapping, patterns across cortical and sub-cortical brain regions [86–89].

At the cellular level, The activation of D_2_/D_3_ dopamine receptors by PPX results in an inhibition of the activity of the PKA and its downstream transduction signaling [90] which leads to a reduction in synaptic strength and eventually promotes long term synaptic depression [91,92]. In the present study PPX treatment led to a reduction in the cellular plasticity marker, Zif268, which plays a major role in synaptic plasticity, especially in the maintenance of long term potentiation (LTP) [55,93]. The dampened expression of Zif268 observed in the PrL and Cg cortex alongside the striatum of HI-PPX rats may reflect a normalization of an otherwise hyperactive goal-directed system in HI rats. In addition, considering the functional antagonism in the control over waiting impulsivity between the NAc Shell and the NAc Core, the latter promoting impulsive action and the former inhibiting it [94–98], the PPX-induced decrease in the mRNA levels of both C-fos and Zif258 in the NAc Core associated with sustained C-fos expression in the Shell of HI rats suggests a PPX-induced ventral striatum-mediated normalization of impulse control at a time HI rats have re-engaged their goal-directed system.

The decreased NAc Core Zif268 mRNA levels associated with optimal impulse control observed here in HI-PPX rats is similar to the profile of LI rats previously established by Besson and colleagues [45]. Together these observations suggest that a decrease in synaptic plasticity in the NAc Core may be a key determinant of behavioral inhibition. In contrast with the Besson study however, no differences were observed here between HI-Veh and LI-Veh rats in the mRNA levels of Zif268 and C-fos in the NAc Core. This apparent discrepancy may be attributable to the difference in the rat strain used in the two studies, namely Sprague-Dawley in the present one and Lister-Hooded in the earlier, a difference in housing conditions (standard versus reversed cycle respectively), a difference in cue-light duration (0.5 vs. 1 s in the Besson and the present study, respectively), which may result in a lower cognitive load in the present study [59,99], as well as differences in training history. Indeed, as compared to the Besson study, rats in the present one were also trained under differential treatment for an additional 12 sessions, an extension of their training history that may have resulted in a lesser recruitment of cellular plasticity processes overall [100].

Taken together, the results of the present study demonstrate a selective anti-impulsive effect of PPX in HI rats under conditions of heightened negative urgency, which is correlated with a specific diminution of the expression of the plasticity marker Zif268 in the NAc Core. These results also identified a specific expression pattern of the cell activity marker, C-fos, maintained by PPX in HI rats in pro-inhibition structures, namely the NAc Shell and DMS and reduced in the pro-impulsive structure, the NAc Core, suggestive of an increased functional recruitment of the mPFC-DMS goal directed system at the expense of the DLS-dependent habit system. These results thereby shed new light on the hitherto accepted pro-impulsive effect of D_2_-like agonists on waiting impulsivity, by revealing a state dependent effect modulated by different corticostriatal circuits. Although further studies are needed to determine whether similar effects are observed in females, these results suggest that targeting D_2_-like receptors may represent a valuable strategy to improve, or normalize, waiting impulsivity in psychiatric disorders in which this dimension of impulsivity is aberrantly exacerbated, such as ADHD or addictions.

## ACKNOWLEGMENTS

The authors want to thank Dr Bizhu He for help with structural equation modeling analysis.

## AUTHOR CONTRIBUTIONS

RM, DB and SC designed experiments with inputs from YMV and SB. RM, MF, YV, TD, and CC performed experiments with help from SB, DB and SC. RM performed the analysis, with help from DB, SC and YC. RM, DB and SC wrote the manuscript with inputs from MF, YMV, YC, TD, CC, SB, MS and PHJ.

## FIGURE LEGENDS

**Supplemental Figure 1:**
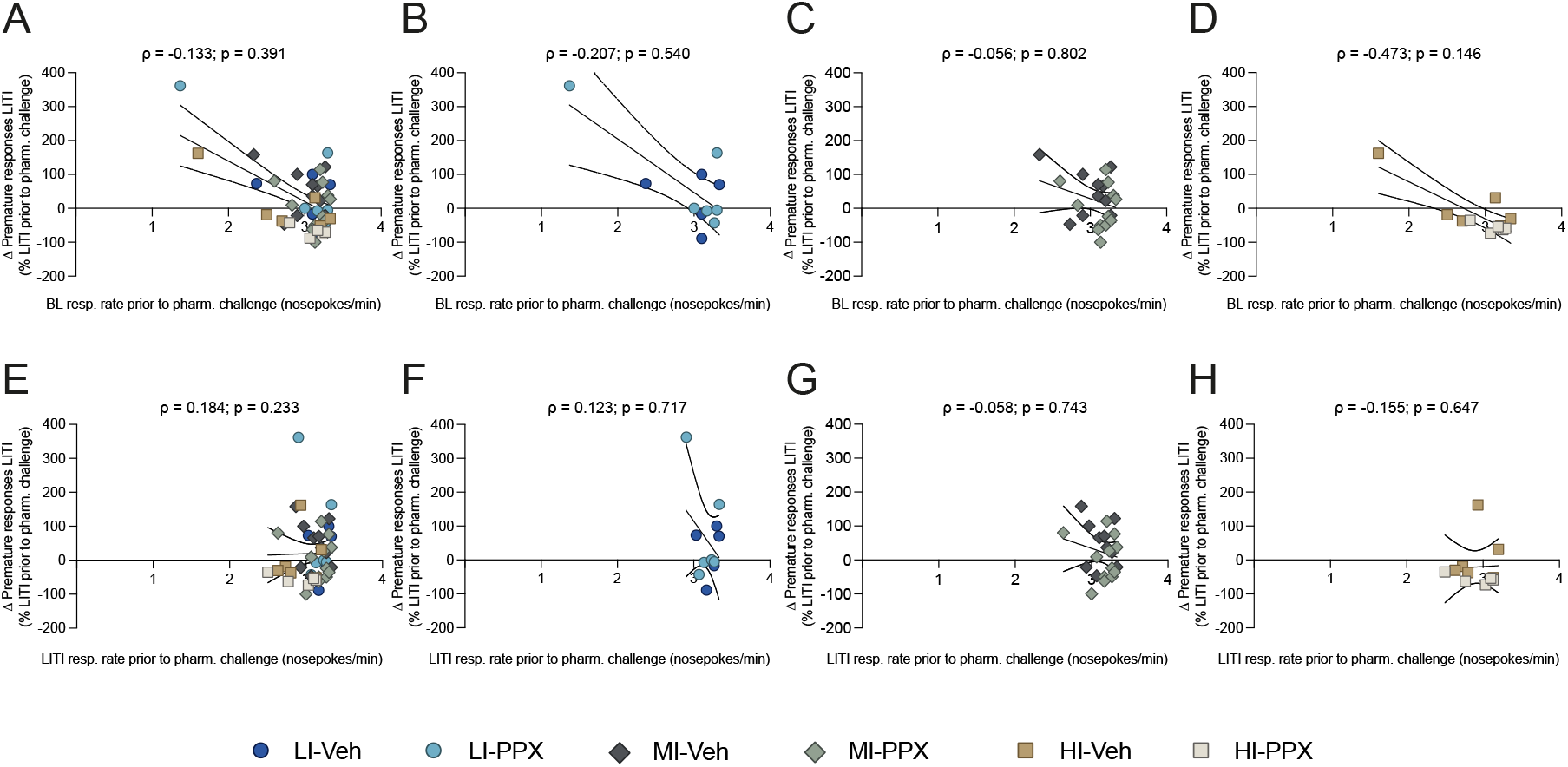
Impulsivity expressed under treatment is not accounted for response rate before treatment. (**A**) Full cohort, (**B**) LI rats, (**C**) MI rats, (**D**) HI rats, premature responding under treatment regressed by response rate during the last baseline (BL) session before treatment. (**E**) Full cohort, (**F**) LI rats, (**G**) MI rats, (**H**) HI rats, premature responding under treatment regressed by response rate during the last long intertrial interval (LITI) session before treatment. LI-Veh n= 5; LI-PPX n = 6; MI-Veh n = 10; MI-PPX n = 12; HI-Veh n = 6; HI-PPX n = 5. LI: low impulsive; MI: moderately impulsive, HI: highly impulsive.

**Supplemental Figure 2:**
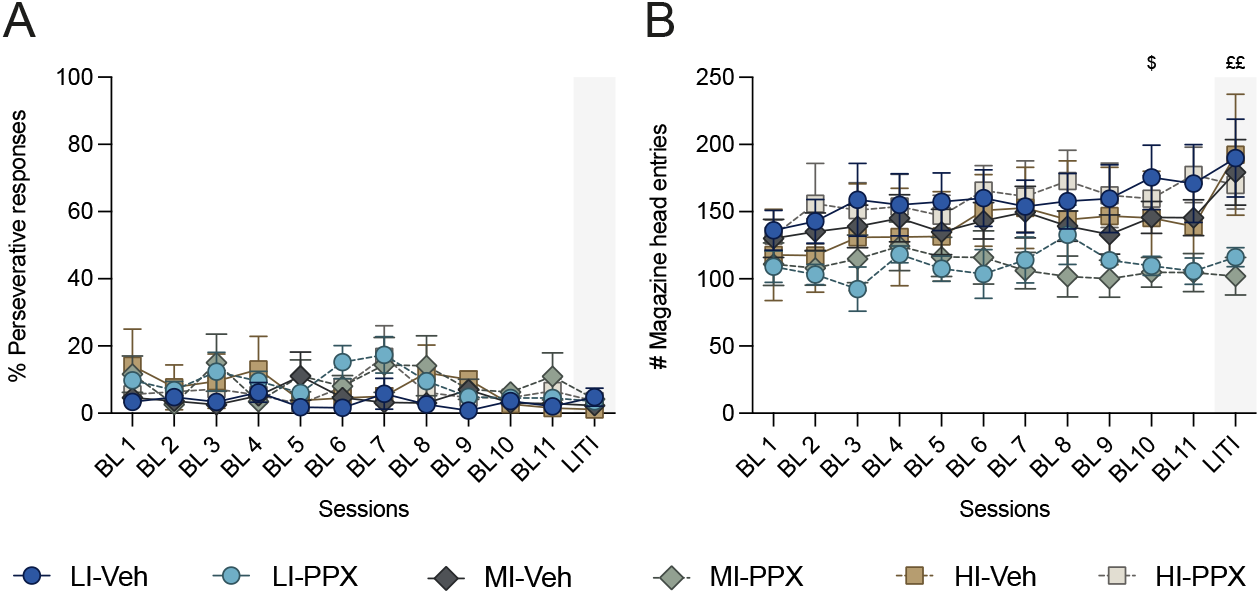
PPX does not induce no perseverative behavior. (**A**) Percentage of perseverative responses made across sessions. (**B**) Number of magazine head entries. LI-Veh n= 5; LI-PPX n = 6; MI-Veh n = 10; MI-PPX n = 12; HI-Veh n = 6; HI-PPX n = 5. BL: baseline; LITI: long intertrial interval; LI: low impulsive; MI: moderately impulsive; HI: higlhy impulsive. Data are shown as means ± SEM. LI-Veh vs. LI-PPX ^$^p<0.05. MI-Veh vs. MI-PPX ^££^p<0.01.

**Supplemental Figure 3:**
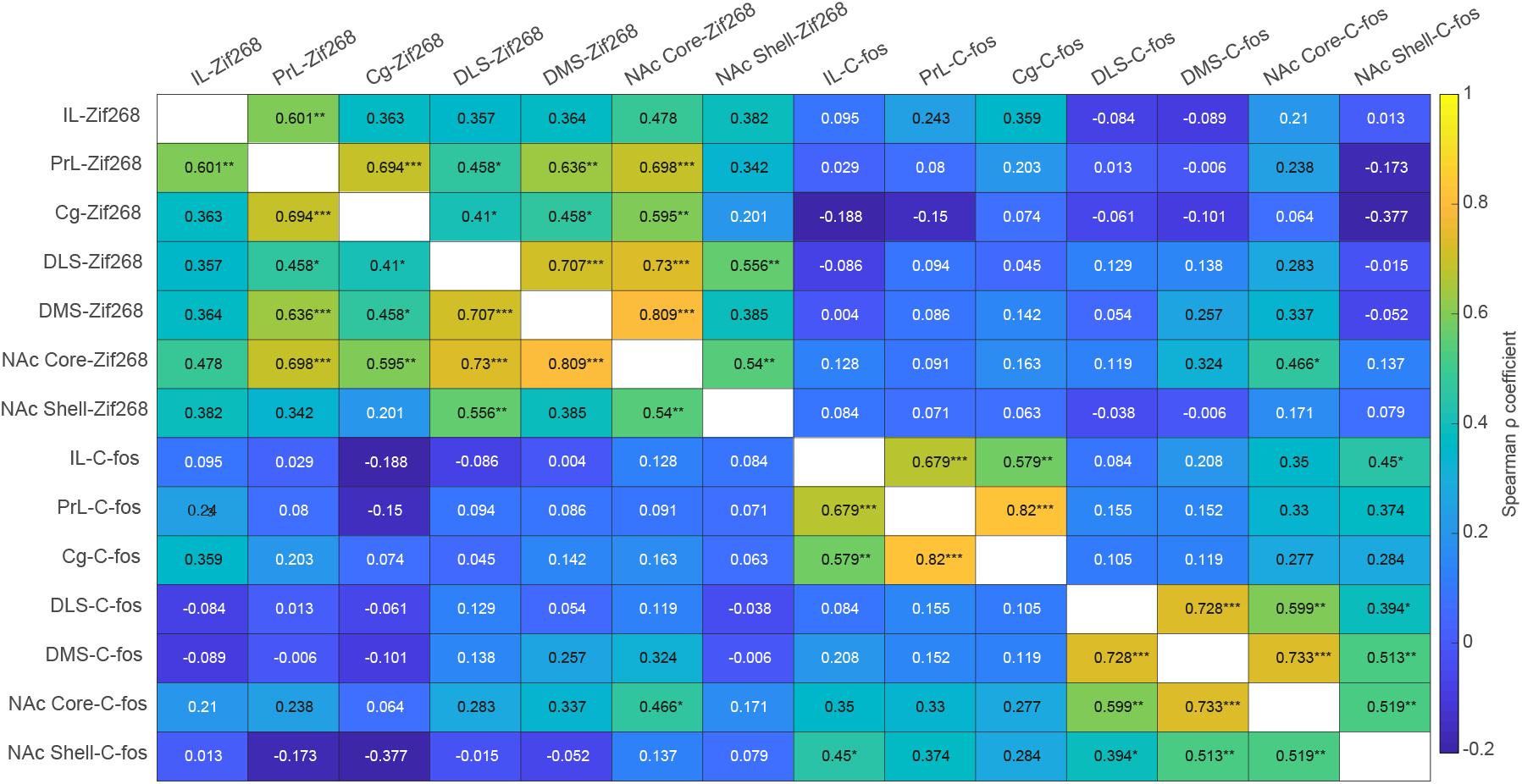
Zif268 and C-fos expression covariance amongst structures of the corticostriatal circuitry. Spearman correlation matrix for C-fos and Zif268 in the seven structures stained for *in situ* hybridization. Color-code represents the ρ value for each covariance, colder colors represent smaller ρ coefficient. LI-Veh n= 5; LI-PPX n = 6; MI-Veh = 8; MI-PPX = 9; HI-Veh n = 6; HI-PPX n = 5. LI: low impulsive; HI: highly impulsive. IL: infralimbic cortex; PrL: prelimbic cortex; Cg: cingulate cortex; DLS: dorsolateral striatum; DMS: dorsomedial striatum; NAc Core: nucleus accumbens core; NAc Shell: nucleus accumbens shell. Significance covariance: ^*^p<0.05; ^**^p<0.01; ^***^p<0.001.

## REFERENCES

1. Houeto J-L, Magnard R, Dalley JW, Belin D, Carnicella S. Trait Impulsivity and Anhedonia: Two Gateways for the Development of Impulse Control Disorders in Parkinson’s Disease? Front Psychiatry. 2016;7.

2. Weintraub D, Koester J, Potenza MN, Siderowf AD, Stacy M, Voon V, et al. Impulse Control Disorders in Parkinson Disease: A Cross-Sectional Study of 3090 Patients. Arch Neurol. 2010;67.

3. Voon V, Sohr M, Lang AE, Potenza MN, Siderowf AD, Whetteckey J, et al. Impulse control disorders in parkinson disease: A multicenter case-control study. Ann Neurol. 2011;69:986–996.

4. Cornelius JR, Tippmann-Peikert M, Slocumb NL, Frerichs CF, Silber MH. Impulse Control Disorders with the use of Dopaminergic Agents in Restless Legs Syndrome: a Case-Control Study. 2010;33:10.

5. Lipford MC, Silber MH. Long-term use of pramipexole in the management of restless legs syndrome. Sleep Med. 2012;13:1280–1285.

6. Voon V, Schoerling A, Wenzel S, Ekanayake V, Reiff J, Trenkwalder C, et al. Frequency of impulse control behaviours associated with dopaminergic therapy in restless legs syndrome. BMC Neurol. 2011;11:117.

7. Holman AJ. Impulse Control Disorder Behaviors Associated with Pramipexole Used to Treat Fibromyalgia. J Gambl Stud. 2009;25:425–431.

8. Bancos I, Nannenga MR, Bostwick JM, Silber MH, Erickson D, Nippoldt TB. Impulse control disorders in patients with dopamine agonist-treated prolactinomas and nonfunctioning pituitary adenomas: a case-control study. Clin Endocrinol (Oxf). 2014;80:863–868.

9. Celik E, Ozkaya HM, Poyraz BC, Saglam T, Kadioglu P. Impulse control disorders in patients with prolactinoma receiving dopamine agonist therapy: a prospective study with 1 year follow-up. Endocrine. 2018;62:692–700.

10. Dogansen SC, Cikrikcili U, Oruk G, Kutbay NO, Tanrikulu S, Hekimsoy Z, et al. Dopamine Agonist-Induced Impulse Control Disorders in Patients With Prolactinoma: A Cross-Sectional Multicenter Study. J Clin Endocrinol Metab. 2019;104:2527–2534.

11. Evans AH, Lees AJ. Dopamine dysregulation syndrome in Parkinson’s disease. Curr Opin Neurol. 2004;17:393–398.

12. Aarsland D, Bronnick K, Alves G, Tysnes OB, Pedersen KF, Ehrt U, et al. The spectrum of neuropsychiatric symptoms in patients with early untreated Parkinson’s disease. J Neurol Neurosurg Psychiatry. 2009;80:928–930.

13. Rabinak CA, Nirenberg MJ. Dopamine Agonist Withdrawal Syndrome in Parkinson Disease. Arch Neurol. 2010;67.

14. Leeman RF, Potenza MN. Similarities and differences between pathological gambling and substance use disorders: a focus on impulsivity and compulsivity. Psychopharmacology (Berl). 2012;219:469–490.

15. Aarts E, Helmich RC, Janssen MJR, Oyen WJG, Bloem BR, Cools R. Aberrant reward processing in Parkinson’s disease is associated with dopamine cell loss. NeuroImage. 2012;59:3339–3346.

16. American Psychiatric Association, American Psychiatric Association, editors. Diagnostic and statistical manual of mental disorders: DSM-5. 5th ed. Washington, D.C: American Psychiatric Association; 2013.

17. Jentsch JD, Ashenhurst JR, Cervantes MC, Groman SM, James AS, Pennington ZT. Dissecting impulsivity and its relationships to drug addictions: Impulsivity subtypes and drug addiction. Ann N Y Acad Sci. 2014:n/a-n/a.

18. Laque A, Wagner GE, Matzeu A, De Ness GL, Kerr TM, Carroll AM, et al. Linking drug and food addiction via compulsive appetite. Br J Pharmacol. 2022;179:2589–2609.

19. Karim R, Chaudhri P. Behavioral Addictions: An Overview. J Psychoactive Drugs. 2012;44:5–17.

20. Yau YHC, Potenza MN. Gambling Disorder and Other Behavioral Addictions: Recognition and Treatment. Harv Rev Psychiatry. 2015;23:134–146.

21. Robbins T, Clark L. Behavioral addictions. Curr Opin Neurobiol. 2015;30:66–72.

22. Perry JL, Carroll ME. The role of impulsive behavior in drug abuse. Psychopharmacology (Berl). 2008;200:1–26.

23. Potenza MN, Balodis IM, Derevensky J, Grant JE, Petry NM, Verdejo-Garcia A, et al. Gambling disorder. Nat Rev Dis Primer. 2019;5:51.

24. Verdejo-Garcia A, Albein-Urios N. Impulsivity traits and neurocognitive mechanisms conferring vulnerability to substance use disorders. Neuropharmacology. 2021;183:108402.

25. Dalley JW, Everitt BJ, Robbins TW. Impulsivity, Compulsivity, and Top-Down Cognitive Control. Neuron. 2011;69:680–694.

26. Evenden JL. Varieties of impulsivity. Psychopharmacology (Berl). 1999;146:348–361.

27. Madden GJ, Johnson PS, Brewer AT, Pinkston JW, Fowler SC. Effects of pramipexole on impulsive choice in male wistar rats. Exp Clin Psychopharmacol. 2010;18:267–276.

28. Koffarnus MN, Newman AH, Grundt P, Rice KC, Woods JH. Effects of selective dopaminergic compounds on a delay-discounting task. Behav Pharmacol. 2011;22:300–311.

29. Martinez E, Pasquereau B, Saga Y, Météreau É, Tremblay L. The Anterior Caudate Nucleus Supports Impulsive Choices Triggered by Pramipexole. Mov Disord. 2020;35:296–305.

30. Johnson PS, Madden GJ, Brewer AT, Pinkston JW, Fowler SC. Effects of acute pramipexole on preference for gambling-like schedules of reinforcement in rats. Psychopharmacology (Berl). 2011;213:11–18.

31. Holtz NA, Tedford SE, Persons AL, Grasso SA, Napier TC. Pharmacologically distinct pramipexole-mediated akinesia vs. risk-taking in a rat model of Parkinson’s disease. Prog Neuropsychopharmacol Biol Psychiatry. 2016;70:77–84.

32. Pes R, Godar SC, Fox AT, Burgeno LM, Strathman HJ, Jarmolowicz DP, et al. Pramipexole enhances disadvantageous decision-making: Lack of relation to changes in phasic dopamine release. Neuropharmacology. 2017;114:77–87.

33. Cole BJ, Robbins TW. Effects of 6-hydroxydopamine lesions of the nucleus accumbens septi on performance of a 5-choice serial reaction time task in rats: Implications for theories of selective attention and arousal. Behav Brain Res. 1989;33:165–179.

34. Robbins T. The 5-choice serial reaction time task: behavioural pharmacology and functional neurochemistry. Psychopharmacology (Berl). 2002;163:362–380.

35. Molander AC, Mar A, Norbury A, Steventon S, Moreno M, Caprioli D, et al. High impulsivity predicting vulnerability to cocaine addiction in rats: some relationship with novelty preference but not novelty reactivity, anxiety or stress. Psychopharmacology (Berl). 2011;215:721–731.

36. Robinson ESJ, Eagle DM, Mar AC, Bari A, Banerjee G, Jiang X, et al. Similar Effects of the Selective Noradrenaline Reuptake Inhibitor Atomoxetine on Three Distinct Forms of Impulsivity in the Rat. Neuropsychopharmacology. 2008;33:1028–1037.

37. Belin-Rauscent A, Daniel M-L, Puaud M, Jupp B, Sawiak S, Howett D, et al. From impulses to maladaptive actions: the insula is a neurobiological gate for the development of compulsive behavior. Mol Psychiatry. 2016;21:491–499.

38. Engeln M, Ansquer S, Dugast E, Bezard E, Belin D, Fernagut P-O. Multi-facetted impulsivity following nigral degeneration and dopamine replacement therapy. Neuropharmacology. 2016;109:69–77.

39. Jiménez-Urbieta H, Gago B, Quiroga-Varela A, Rodríguez-Chinchilla T, Merino-Galán L, Oregi A, et al. Pramipexole-induced impulsivity in mildparkinsonian rats: a model of impulse control disorders in Parkinson’s disease. Neurobiol Aging. 2019;75:126–135.

40. Belin D, Mar AC, Dalley JW, Robbins TW, Everitt BJ. High Impulsivity Predicts the Switch to Compulsive Cocaine-Taking. Science. 2008;320:1352–1355.

41. Ersche KD, Barnes A, Jones PS, Morein-Zamir S, Robbins TW, Bullmore ET. Abnormal structure of frontostriatal brain systems is associated with aspects of impulsivity and compulsivity in cocaine dependence. Brain. 2011;134:2013–2024.

42. Ansquer S, Belin-Rauscent A, Dugast E, Duran T, Benatru I, Mar AC, et al. Atomoxetine Decreases Vulnerability to Develop Compulsivity in High Impulsive Rats. Biol Psychiatry. 2014;75:825–832.

43. Jones JA, Belin-Rauscent A, Jupp B, Fouyssac M, Sawiak SJ, Zuhlsdorff K, et al. Neurobehavioral precursors of compulsive cocaine-seeking in dual fronto-striatal circuits. Neuroscience; 2022.

44. Eagle DM, Baunez C. Is there an inhibitory-response-control system in the rat? Evidence from anatomical and pharmacological studies of behavioral inhibition. Neurosci Biobehav Rev. 2010;34:50–72.

45. Besson M, Pelloux Y, Dilleen R, Theobald DE, Lyon A, Belin-Rauscent A, et al. Cocaine Modulation of Frontostriatal Expression of Zif268, D2, and 5-HT2c Receptors in High and Low Impulsive Rats. Neuropsychopharmacology. 2013;38:1963–1973.

46. Murray JE, Dilleen R, Pelloux Y, Economidou D, Dalley JW, Belin D, et al. Increased Impulsivity Retards the Transition to Dorsolateral Striatal Dopamine Control of Cocaine Seeking. Biol Psychiatry. 2014;76:15–22.

47. Giuliano C, Marti-Prats L, Domi A, Puaud M, Pena-Oliver Y, McKenzie C, et al. The development of compulsive coping behaviors depends on the engagement of dorsolateral striatum dopamine-dependent mechanisms. Neuroscience; 2022.

48. Hope B, Kosofsky B, Hyman SE, Nestler EJ. Regulation of immediate early gene expression and AP-1 binding in the rat nucleus accumbens by chronic cocaine. Proc Natl Acad Sci. 1992;89:5764–5768.

49. Crombag HS, Jedynak JP, Redmond K, Robinson TE, Hope BT. Locomotor sensitization to cocaine is associated with increased Fos expression in the accumbens, but not in the caudate. Behav Brain Res. 2002;136:455–462.

50. Cruz FC, Javier Rubio F, Hope BT. Using c-fos to study neuronal ensembles in corticostriatal circuitry of addiction. Brain Res. 2015;1628:157–173.

51. Thomas KL, Arroyo M, Everitt BJ. Induction of the learning and plasticity-associated gene Zif268 following exposure to a discrete cocaine-associated stimulus: Cocaine CS induced Zif268 expression. Eur J Neurosci. 2003;17:1964–1972.

52. Valjent E. Plasticity-Associated Gene Krox24/Zif268 Is Required for Long-Lasting Behavioral Effects of Cocaine. J Neurosci. 2006;26:4956–4960.

53. Hearing MC, See RE, McGinty JF. Relapse to cocaine-seeking increases activity-regulated gene expression differentially in the striatum and cerebral cortex of rats following short or long periods of abstinence. Brain Struct Funct. 2008;213:215–227.

54. Unal CT, Beverley JA, Willuhn I, Steiner H. Long-lasting dysregulation of gene expression in corticostriatal circuits after repeated cocaine treatment in adult rats: effects on zif 268 and homer 1a. Eur J Neurosci. 2009;29:1615–1626.

55. Veyrac A, Besnard A, Caboche J, Davis S, Laroche S. The Transcription Factor Zif268/Egr1, Brain Plasticity, and Memory. Prog. Mol. Biol. Transl. Sci., vol. 122, Elsevier; 2014. p. 89–129.

56. Carli M, Robbins TW, Evenden JL, Everitt BJ. Effects of lesions to ascending noradrenergic neurones on performance of a 5-choice serial reaction task in rats; implications for theories of dorsal noradrenergic bundle function based on selective attention and arousal. Behav Brain Res. 1983;9:361–380.

57. Bari A, Dalley JW, Robbins TW. The application of the 5-choice serial reaction time task for the assessment of visual attentional processes and impulse control in rats. Nat Protoc. 2008;3:759–767.

58. Auclair AL, Besnard J, Newman-Tancredi A, Depoortère R. The five choice serial reaction time task: Comparison between Sprague–Dawley and Long–Evans rats on acquisition of task, and sensitivity to phencyclidine. Pharmacol Biochem Behav. 2009;92:363–369.

59. Turner KM, Peak J, Burne THJ. Measuring Attention in Rodents: Comparison of a Modified Signal Detection Task and the 5-Choice Serial Reaction Time Task. Front Behav Neurosci. 2016;9.

60. Turner KM, Young JW, McGrath JJ, Eyles DW, Burne THJ. Cognitive performance and response inhibition in developmentally vitamin D (DVD)-deficient rats. Behav Brain Res. 2013;242:47–53.

61. Favier M, Duran T, Carcenac C, Drui G, Savasta M, Carnicella S. Pramipexole reverses Parkinson’s disease-related motivational deficits in rats: Pramipexole and Motivation. Mov Disord. 2014;29:912–920.

62. Magnard R, Vachez Y, Carcenac C, Krack P, David O, Savasta M, et al. What can rodent models tell us about apathy and associated neuropsychiatric symptoms in Parkinson’s disease? Transl Psychiatry. 2016;6:e753–e753.

63. Chang W, Breier MR, Yang A, Swerdlow NR. Disparate effects of pramipexole on locomotor activity and sensorimotor gating in Sprague–Dawley rats. Pharmacol Biochem Behav. 2011;99:634–638.

64. Maj J, Rogóż Z, Skuza G, Kołodziejczyk K. The behavioural effects of pramipexole, a novel dopamine receptor agonist. Eur J Pharmacol. 1997;324:31–37.

65. Belin D, Deroche-Gamonet V, Jaber M. Cocaine-induced sensitization is associated with altered dynamics of transcriptional responses of the dopamine transporter, tyrosine hydroxylase, and dopamine D2 receptors in C57Bl/6J mice. Psychopharmacology (Berl). 2007;193:567–578.

66. Fouyssac M, Peña-Oliver Y, Puaud M, Lim NTY, Giuliano C, Everitt BJ, et al. Negative Urgency Exacerbates Relapse to Cocaine Seeking After Abstinence. Biol Psychiatry. 2022;91:1051–1060.

67. Schneider CA, Rasband WS, Eliceiri KW. NIH Image to ImageJ: 25 years of image analysis. Nat Methods. 2012;9:671–675.

68. Paxinos G, Watson C. The rat brain in stereotaxic coodinates. San diego: Elsevier Academic Press. 1998.

69. Levine TR, Hullett CR. Eta Squared, Partial Eta Squared, and Misreporting of Effect Size in Communication Research. Hum Commun Res. 2002;28:612–625.

70. Magnard R, Vachez Y, Carcenac C, Boulet S, Houeto J-L, Savasta M, et al. Nigrostriatal Dopaminergic Denervation Does Not Promote Impulsive Choice in the Rat: Implication for Impulse Control Disorders in Parkinson’s Disease. Front Behav Neurosci. 2018;12:312.

71. Baron RM, Kenny DA. The Moderator-Mediator Variable Distinction in Social Psychological Research: Conceptual, Strategic, and Statistical Considerations. J Pers Soc Psychol. 1986;51:1173–1182.

72. Ditlevsen S, Christensen U, Lynch J, Damsgaard MT, Keiding N. The Mediation Proportion: A Structural Equation Approach for Estimating the Proportion of Exposure Effect on Outcome Explained by an Intermediate Variable. Epidemiology. 2005;16:114–120.

73. Brown VJ, Robbins TW. Simple and choice reaction time performance following unilateral striatal dopamine depletion in the rat: impaired motor readiness but preserved response preparation. Brain. 1991;114:513–525.

74. Tomlinson A, Grayson B, Marsh S, Harte MK, Barnes SA, Marshall KM, et al. Pay attention to impulsivity: Modelling low attentive and high impulsive subtypes of adult ADHD in the 5-choice continuous performance task (5C-CPT) in female rats. Eur Neuropsychopharmacol. 2014;24:1371–1380.

75. Caprioli D, Jupp B, Hong YT, Sawiak SJ, Ferrari V, Wharton L, et al. Dissociable Rate-Dependent Effects of Oral Methylphenidate on Impulsivity and D2/3 Receptor Availability in the Striatum. J Neurosci. 2015;35:3747–3755.

76. Higgins GA, Silenieks LB, MacMillan C, Thevarkunnel S, Parachikova AI, Mombereau C, et al. Characterization of Amphetamine, Methylphenidate, Nicotine, and Atomoxetine on Measures of Attention, Impulsive Action, and Motivation in the Rat: Implications for Translational Research. Front Pharmacol. 2020;11:427.

77. Fernando ABP, Economidou D, Theobald DE, Zou M-F, Newman AH, Spoelder M, et al. Modulation of high impulsivity and attentional performance in rats by selective direct and indirect dopaminergic and noradrenergic receptor agonists. Psychopharmacology (Berl). 2012;219:341–352.

78. Schippers MC, Bruinsma B, Gaastra M, Mesman TI, Denys D, De Vries TJ, et al. Deep Brain Stimulation of the Nucleus Accumbens Core Affects Trait Impulsivity in a Baseline-Dependent Manner. Front Behav Neurosci. 2017;11.

79. Dalley JW, Fryer TD, Brichard L, Robinson ESJ, Theobald DEH, Lääne K, et al. Nucleus Accumbens D2/3 Receptors Predict Trait Impulsivity and Cocaine Reinforcement. 2007;315:5.

80. Zeeb FD, Soko AD, Ji X, Fletcher PJ. Low Impulsive Action, but not Impulsive Choice, Predicts Greater Conditioned Reinforcer Salience and Augmented Nucleus Accumbens Dopamine Release. Neuropsychopharmacology. 2016;41:2091–2100.

81. Fitzpatrick CM, Maric VS, Bate ST, Andreasen JT. Influence of intertrial interval on basal and drug-induced impulsive action in the 5-choice serial reaction time task: Effects of damphetamine and (±)-2,5-dimethoxy-4-iodoamphetamine (DOI). Neurosci Lett. 2018;662:351–355.

82. Toschi C, El-Sayed Hervig M, Burghi T, Sell T, Lycas MD, Moazen P, et al. Dissociating reward sensitivity and negative urgency effects on impulsivity in the five-choice serial reaction time task. Brain Neurosci Adv. 2022;6:239821282211022.

83. Yin HH, Knowlton BJ, Balleine BW. Lesions of dorsolateral striatum preserve outcome expectancy but disrupt habit formation in instrumental learning. Eur J Neurosci. 2004;19:181–189.

84. Faure A. Lesion to the Nigrostriatal Dopamine System Disrupts Stimulus-Response Habit Formation. J Neurosci. 2005;25:2771–2780.

85. Terra H, Bruinsma B, de Kloet SF, van der Roest M, Pattij T, Mansvelder HD. Prefrontal Cortical Projection Neurons Targeting Dorsomedial Striatum Control Behavioral Inhibition. Curr Biol. 2020;30:4188–4200.e5.

86. Brockett AT, Tennyson SS, deBettencourt CA, Gaye F, Roesch MR. Anterior cingulate cortex is necessary for adaptation of action plans. Proc Natl Acad Sci. 2020;117:6196–6204.

87. Covington HE, Lobo MK, Maze I, Vialou V, Hyman JM, Zaman S, et al. Antidepressant Effect of Optogenetic Stimulation of the Medial Prefrontal Cortex. J Neurosci. 2010;30:16082–16090.

88. Guzowski JF, Setlow B, Wagner EK, McGaugh JL. Experience-Dependent Gene Expression in the Rat Hippocampus after Spatial Learning: A Comparison of the Immediate-Early Genes Arc, c-fos, and zif268. J Neurosci. 2001;21:5089–5098.

89. Brami-Cherrier K, Valjent E, Hervé D, Darragh J, Corvol J-C, Pages C, et al. Parsing Molecular and Behavioral Effects of Cocaine in Mitogen- and Stress-Activated Protein Kinase-1-Deficient Mice. J Neurosci. 2005;25:11444–11454.

90. Neve KA, Seamans JK, Trantham-Davidson H. Dopamine Receptor Signaling. J Recept Signal Transduct. 2004;24:165–205.

91. Sun X, Zhao Y, Wolf ME. Dopamine Receptor Stimulation Modulates AMPA Receptor Synaptic Insertion in Prefrontal Cortex Neurons. J Neurosci. 2005;25:7342–7351.

92. Shen W, Flajolet M, Greengard P, Surmeier DJ. Dichotomous Dopaminergic Control of Striatal Synaptic Plasticity. Science. 2008;321:848–851.

93. Jones MW, Errington ML, French PJ, Fine A, Bliss TVP, Garel S, et al. A requirement for the immediate early gene Zif268 in the expression of late LTP and long-term memories. Nat Neurosci. 2001;4:289–296.

94. Sesia T, Temel Y, Lim LW, Blokland A, Steinbusch HWM, Visser-Vandewalle V. Deep brain stimulation of the nucleus accumbens core and shell: Opposite effects on impulsive action. Exp Neurol. 2008;214:135–139.

95. Sesia T, Bulthuis V, Tan S, Lim LW, Vlamings R, Blokland A, et al. Deep brain stimulation of the nucleus accumbens shell increases impulsive behavior and tissue levels of dopamine and serotonin. Exp Neurol. 2010;225:302–309.

96. Moreno M, Economidou D, Mar AC, López-Granero C, Caprioli D, Theobald DE, et al. Divergent effects of D2/3 receptor activation in the nucleus accumbens core and shell on impulsivity and locomotor activity in high and low impulsive rats. Psychopharmacology (Berl). 2013;228:19–30.

97. Feja M, Hayn L, Koch M. Nucleus accumbens core and shell inactivation differentially affects impulsive behaviours in rats. Prog Neuropsychopharmacol Biol Psychiatry. 2014;54:31–42.

98. Feja M, Koch M. Frontostriatal systems comprising connections between ventral medial prefrontal cortex and nucleus accumbens subregions differentially regulate motor impulse control in rats. Psychopharmacology (Berl). 2015;232:1291–1302.

99. Caballero-Puntiverio M, Fitzpatrick CM, Woldbye DP, Andreasen JT. Effects of amphetamine and methylphenidate on attentional performance and impulsivity in the mouse 5-Choice Serial Reaction Time Task. J Psychopharmacol (Oxf). 2017;31:272–283.

100. Hernandez PJ, Schiltz CA, Kelley AE. Dynamic shifts in corticostriatal expression patterns of the immediate early genes Homer 1a and Zif268 during early and late phases of instrumental training. Learn Mem. 2006;13:599–608.

